# Identification of differential polypharmacology between the PARP inhibitor rucaparib and its major metabolite

**DOI:** 10.1101/2022.11.22.517505

**Authors:** Huabin Hu, Carme Serra, Amadeu Llebaria, Albert A. Antolin

**Author notes:** Correspondence (A.A.A.), (A.L.).

## Abstract

The (poly)pharmacology of drug metabolites is seldom comprehensively characterized in drug discovery and development. However, some drug metabolites can reach high plasma concentrations and display relevant in vivo activity, which can be distinct from its parent drug. Here, we use computational and experimental methods to comprehensively characterise the kinase polypharmacology of M324, the major metabolite of the FDA-approved PARP inhibitor rucaparib. We experimentally demonstrate that M324 displays a distinct in vitro kinome profile from its parent drug, characterized by potent in vitro inhibition of GSK3A and PLK2 at clinically-relevant concentrations. These confirmed kinase activities of M324 could have potential implications for the efficacy and safety of rucaparib and therefore warrant further clinical investigation. The study reported here highlights the importance of thoroughly characterizing the activity of significant drug metabolites to better understanding drug responses in the clinic and maximally exploit the current drug arsenal in personalized and precision medicine.

## Introduction

Small molecule drugs administered orally are normally metabolized through various proteins (e.g., cytochrome P450 (CYP450) enzymes in the liver) that catalyse distinct chemical modifications aimed at increasing aqueous solubility to facilitate excretion (Fura, 2006; Guengerich, 2001; Zhang and Tang, 2018). Accordingly, drug metabolites often show lower cellular permeability and plasma concentration than their parent drugs, preventing them from displaying relevant *in vivo* pharmacological activity (Fura, 2006; Zhang and Tang, 2018). Nonetheless, certain drug metabolites are present at high plasma concentrations and can permeate inside human cells or interact with membrane receptors to display cellular activity (Fura, 2006). In addition, the chemical modifications performed on drug metabolites can profoundly alter their interaction with human proteins (Fura, 2006; Thompson et al., 2016; Walther et al., 2017). In these cases, metabolites can have a distinct biological activity than their parent drugs, which can have implications for both *in vivo* efficacy and safety (Baillie and Rettie, 2011; Bender et al., 2004; Thompson et al., 2016). Moreover, the simultaneous presence of active metabolites and parent drugs in the body could even synergize to produce either positive or detrimental effects (Fura, 2006). Unfortunately, drug metabolites are rarely characterized comprehensively in preclinical models, are often not commercially available to facilitate testing by the wide scientific community, and their protein binding activity is often overlooked in drug development, particularly beyond the known targets of their parent drug. As any other small molecule, drug metabolites can interact with multiple protein targets simultaneously, a behaviour termed polypharmacology (Anighoro et al., 2014; Bolognesi, 2019; Hopkins, 2008; Kabir and Muth, 2022). Therefore, systematically characterizing the polypharmacology of cell-permeable drug metabolites that are present at high concentrations could help clarify the unexplained clinical activity of drugs, open new repurposing opportunities and help tailor drugs to patients.

Poly(ADP-ribose) polymerase-1 (PARP1) inhibitors are an established class of targeted, small-molecule drugs approved for various types of cancers displaying alterations in DNA repair genes (Mateo et al., 2019; Rose et al., 2020; Sinha et al., 2021). Thus far, four PARP1 inhibitors (olaparib, rucaparib, talazoparib, and niraparib) have been approved by the FDA (Antolin et al., 2020). When analyzing the chemical structures of PARP1 drugs it becomes apparent that all approved PARP1 inhibitors share a common benzamide substructure (either a primary amide or a cyclic lactam, **Figure 1**), which mimics the nicotinamide moiety of PARP1’s substrate NAD^+^ to form a hydrogen bond network with critical amino acid residues in the binding site (Ser904 and Gly863), as shown in **Figure 1** (bottom). Moreover, the aromatic ring system where the benzamide group is attached forms a π-π stacking interaction with residue Tyr907 (**Figure 1**) (Ferraris, 2010). Given these conserved interactions, FDA-approved PARP inhibitors share a similar profile against PARP enzyme family members, displaying low selectivity between PARP isoforms 1-3, and showing modest activity against other PARP family members (Antolin et al., 2020; Thorsell et al., 2017). Accordingly, current PARP inhibitors share many side effects (Antolin et al., 2020; LaFargue et al., 2019; Sandhu et al., 2022) which can be attributed to a comparable inhibitory interaction with PARP enzymes. However, it has also been reported that there are some unique side effects specific to each PARP inhibitor that cannot be explained by their similar selectivity profiles against PARP enzymes (Antolin et al., 2020; LaFargue et al., 2019).

**Figure 1.**
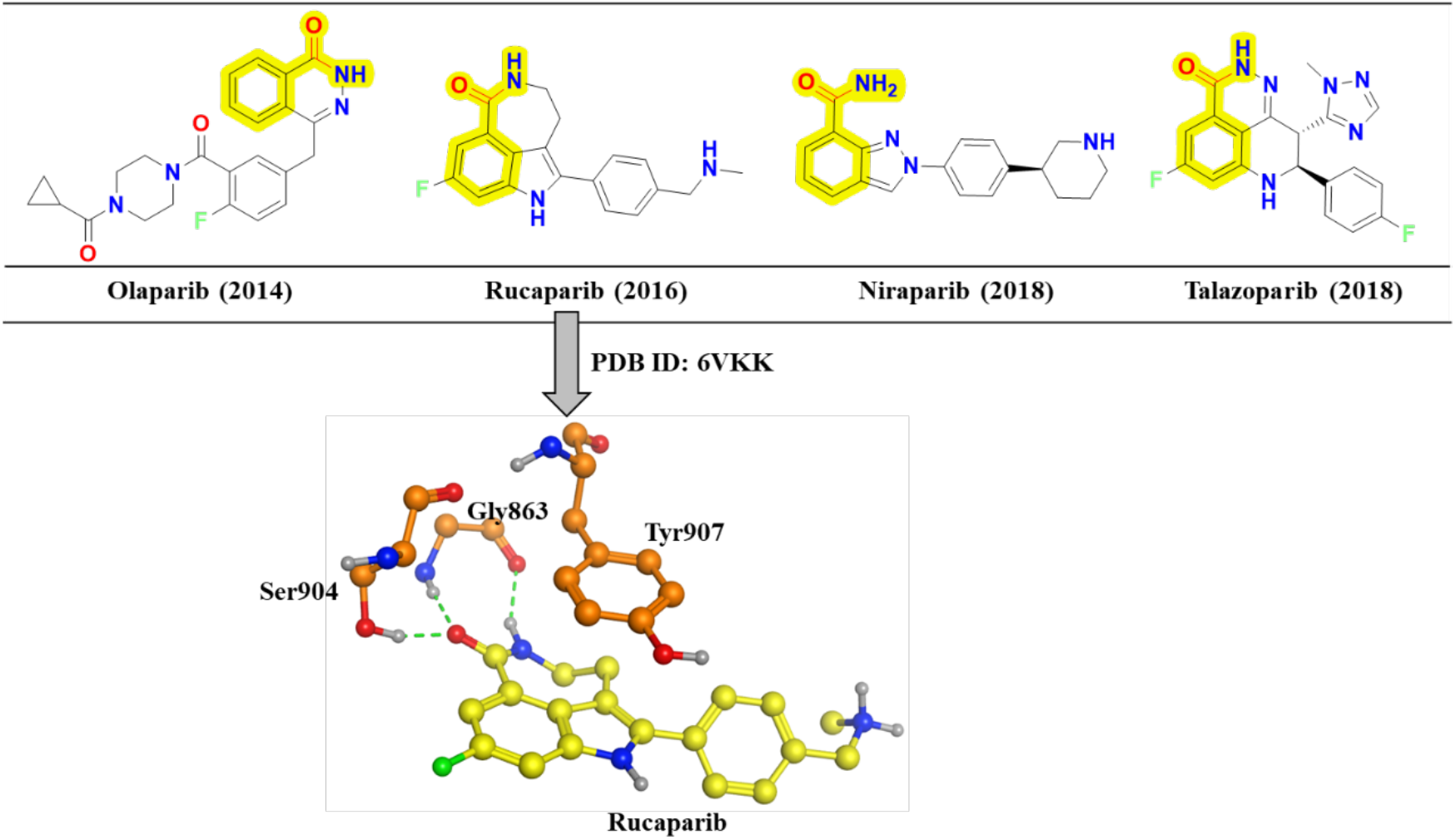
FDA approved PARP1 inhibitors and key molecular interactions. The chemical structures of the four PARP1 inhibitors approved by the FDA are displayed in the upper panel. The substructure (benzamide) mimicking the nicotinamide moiety of the PARP1 substrate NAD^+^ is highlighted in yellow on each chemical structure. The year of the first FDA approval is shown in brackets. The lower panel exemplifies the key conserved interactions between PARP1 and its inhibitors using rucaparib as an example. For clarity, only key interactions with the core scaffold of rucaparib are depicted. The carbons of key PARP1 residues and rucaparib are coloured orange and yellow, respectively.

Despite their conserved interactions, all FDA-approved PARP1 drugs have a unique chemical structure with a different core and substituents (**Figure 1**). Our group, alongside others, has recently used computational and experimental methods to uncover that the unique chemical structure of each approved PARP inhibitor translates into different polypharmacology profiles across the human kinome (Antolin et al., 2020; Knezevic et al., 2016). Whilst olaparib and talazoparib are unlikely to modulate kinase activity at relevant concentrations (10 µM), niraparib and rucaparib showed significant kinase polypharmacology (Antolin et al., 2020). In particular, rucaparib exhibited submicromolar *in vitro* activity against DYRK1B, CDK16, and PIM3, whereas niraparib potently inhibited DYRK1A and DYRK1B (Antolin et al., 2020). The systematic kinome-wide polypharmacology profiling enabled us to link some of the differential side effects observed between PARP inhibitors to their different kinase off-target activities. For example, unique PIM3 inhibition by rucaparib could explain the increased cholesterol levels observed in some patients taking this drug, which have not been observed with other PARP inhibitors (Antolin et al., 2020). However, many differential side effects between PARP inhibitors remain unexplained, such as the increased number of cardiac adverse drug reactions, including arrhythmias, observed in patients taking rucaparib that are not observed with olaparib or talazoparib (Sandhu et al., 2022).

In general, rucaparib has a low metabolic turnover rate in human liver microsomes and *in vitro* metabolite profiling shows that rucaparib is metabolized predominantly by CYP2D6 and, to a lesser extent, by CYP1A2 and CYP3A4 (https://www.accessdata.fda.gov/drugsatfda_docs/nda/2016/209115Orig1s000MultiDisciplineR.pdf). The phase I oxidative deamination reaction catalyzed by CYP450-related enzymes results in the conversion of rucaparib to its major metabolite M324, a carboxylic acid, (**Figure 2a**) which can be detected in several species, including mouse, rat, dog, monkey, and human (Liao et al., 2020; Murray et al., 2014). In animal models (Capan-1 tumour-bearing mice), pharmacokinetic analysis demonstrated that M324 can reach higher concentrations in plasma than the parent drug but with lower concentration inside the tumour cells. Still, M324 was able to reach single digit micromolar concentrations inside mice tumour cells and was still detectable two days after oral administration (Murray et al., 2014). In humans, after giving a single oral dose of 600 mg [^14^C]-rucaparib to patients with advanced solid tumours, M324 was the major detectable metabolite in plasma with a concentration up to ~ 40% of rucaparib’s concentration (Liao et al., 2020). Although a study reported that M324 showed limited cellular enzyme activity against PARP1 in intact human cells (Murray et al., 2014) and it has been reported to be “at least 30 fold less potent than rucaparib against PARP1-3” (https://www.ema.europa.eu/en/documents/product-information/rubraca-epar-product-information_en.pdf), the micromolar concentrations that M324 can reach inside mice tumours demonstrates its capacity to permeate inside cells. Therefore, its potential pharmacological activity, particularly beyond the PARP enzyme family, warrants further investigation and could help explain the unique clinical activity of rucaparib and facilitate its precise and personalized use.

**Figure 2.**
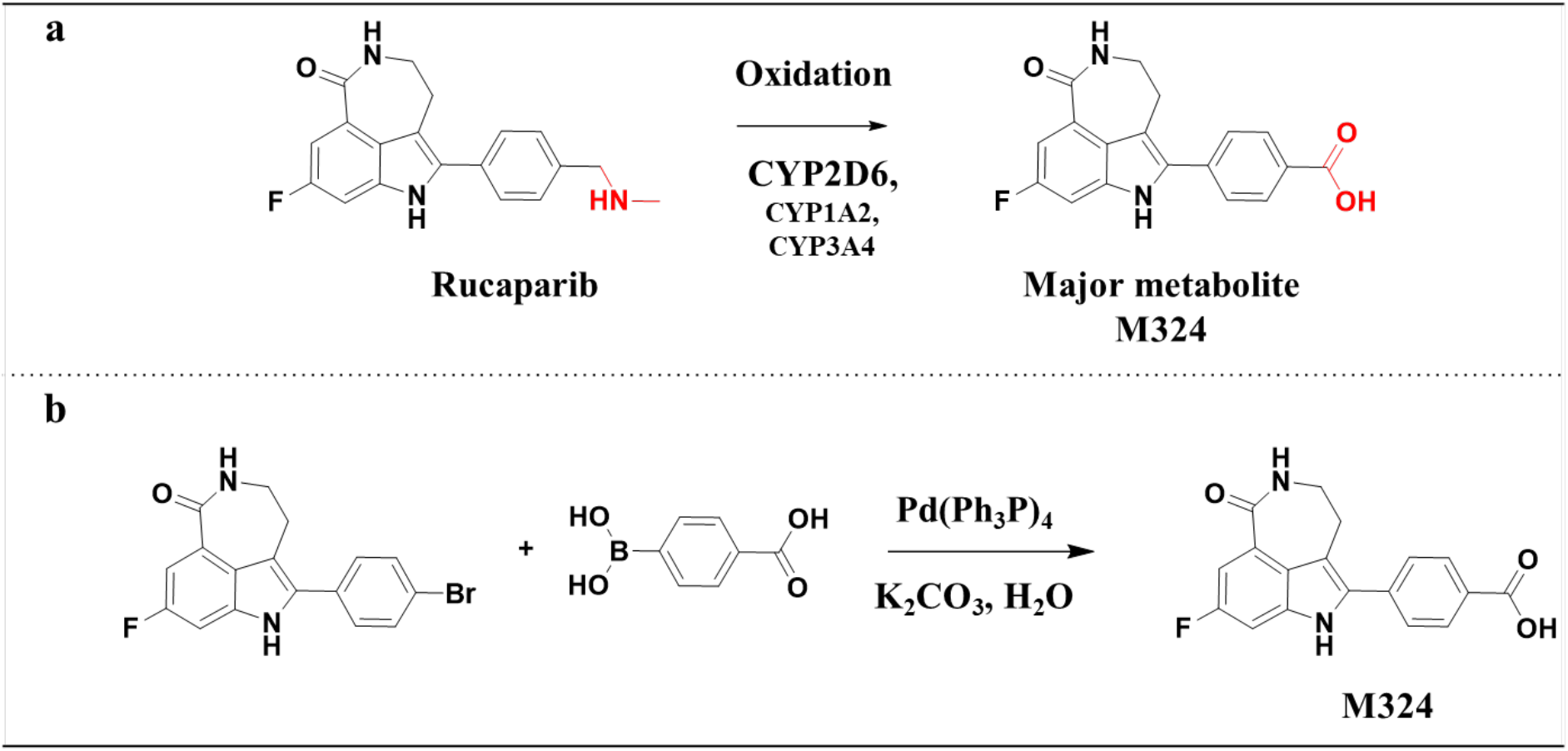
Biotransformation of rucaparib to its major metabolite and corresponding synthesis scheme. **(a)** Cytochrome P450-mediated oxidative deamination of rucaparib, which yields the major product (M324). The structural modification showing the change from an amine to a carboxylic acid is coloured red. **(b)** Synthesis of the metabolite M324 using a Suzuki cross-coupling reaction.

In this study, we have comprehensively characterized the kinase off-target profile of the M324 metabolite. We firstly characterized its kinome profile using computational approaches followed by experimental validation. Our computational results confirmed our hypothesis that M324 and its parent drug have differential kinase polypharmacology landscapes that could have implications for the clinical efficacy and safety of rucaparib. We then synthesized M324 via one-step Suzuki cross-coupling reaction and experimentally tested the predicted kinase activities. The detailed results are reported here.

## Results

### Chemical synthesis of the major metabolite M324

Most drugs are readily available for purchase from chemical vendors. Unfortunately, most drug metabolites are absent from vendor catalogues, limiting their experimental study and characterization by the wider scientific community. Accordingly, we first synthesized this metabolite via one-step Suzuki cross-coupling reaction starting with two available reagents (2-bromo-8-fluoro-4,5-dihydro-1H-azepino[5,4,3-cd]indol-6(3H)-one and 4-boronobenzoic acid), as illustrated in **Figure 2b**. The resulting product was purified and obtained as a greenish solid (71% yield). The chemical structure of the compound has been confirmed by NMR and was ≥ 95% pure by HPLC/MS (**Figures S1-S2**).

### Kinase polpyharmacological profiles predicted using different computational approaches

Various computational methodologies have been developed for polypharmacology prediction based on integrating public bioactivity data from different sources (Cereto-Massague et al., 2015; Lavecchia and Cerchia, 2016). In this study, we employed four different *in silico* approaches, covering both ligand- and structure-based methods, to predict potential kinase off-targets of rucaparib and M324, namely: (1) the six independent ligand-based methods integrated in the CLARITY platform developed by Chemotargets (Vidal et al., 2011); (2) the similarity ensemble approach (SEA) (Lounkine et al., 2012), a web-based tool to conduct target prediction based on chemical similarity; (3) the polypharmacology browser PPB2 incorporating different models (e.g. fingerprint comparison, machine learning or deep learning) to predict potential targets for compounds (Awale and Reymond, 2019); (4) GalaxySagittarius software which integrates ligand- and structure-based approaches to derive target hypothesis for a query compound (Yang et al., 2020). The results of the computational methods concerning only human protein kinases are summarized in **Table 1** (and **Tables S1–S4**).

**Table 1.** *In silico* and *in vitro* kinome profiling statistics for rucaparib and M324. The number of kinases predicted using the four different computational methods used (see **STAR methods**) and the number of real kinase hits confirmed by experimental testing (≤ 50% kinase activity at 10 µM threshold) are shown for both rucaparib and M324. The number in parenthesis represents the true predictions confirmed by experimental testing. Rucaparib was predicted and experimentally confirmed to inhibit a higher number of kinases, however, all the computational methods failed to identify several of the experimental hits and predicted a significant number of false positives.

| Compound name | Computational prediction for kinase targets (**true**) | | | | | Experimental testing (enzyme activity $\leq 50\%$) |
| --- | --- | --- | --- | --- | --- | --- |
|  | CLARITY | SEA | PPB2 | GalaxySagittarius | Total |  |
| Rucaparib | 7 (2) | 8 (2) | 13 (2) | 66 (9) | 75 (9) | 50 |
| M324 | 7 (2) | 5 (2) | 19 (3) | 39 (6) | 56 (9) | 28 |

In total, 75 and 56 kinases were predicted to be potential kinase off-targets for rucaparib and its major metabolite, respectively. Most of the kinase predictions for both compounds are derived from GalaxySagittarius (66 kinases predicted for rucaparib and 39 for M324), followed by PPB2 with 13 and 19 kinases for the drug and the metabolite, respectively. CLARITY and SEA predicted fewer than 10 kinases for both compounds (**Table 1**; **Tables S1–S4**). **Figure 3** provides a network overview of the computational results which connects the predicted kinase targets with either rucaparib or M324. As expected, these two compounds with highly structural similarity have many overlapping predicted kinases (38 green nodes, **Figure 3**). However, there are also many kinases that are only predicted to bind to the drug or the metabolite: 37 kinases (blue nodes, **Figure 3**) are exclusively predicted for rucaparib and 18 kinases (yellow nodes) for M324. From the predicted compound-kinase network, we hypothesized that M324 could be modulating several unique kinases from those modulated by rucaparib. This different polypharmacology profile could help us rationalize the unique rucaparib-associated side effects observed in the clinic as compared to other PARP inhibitors.

**Figure 3.**
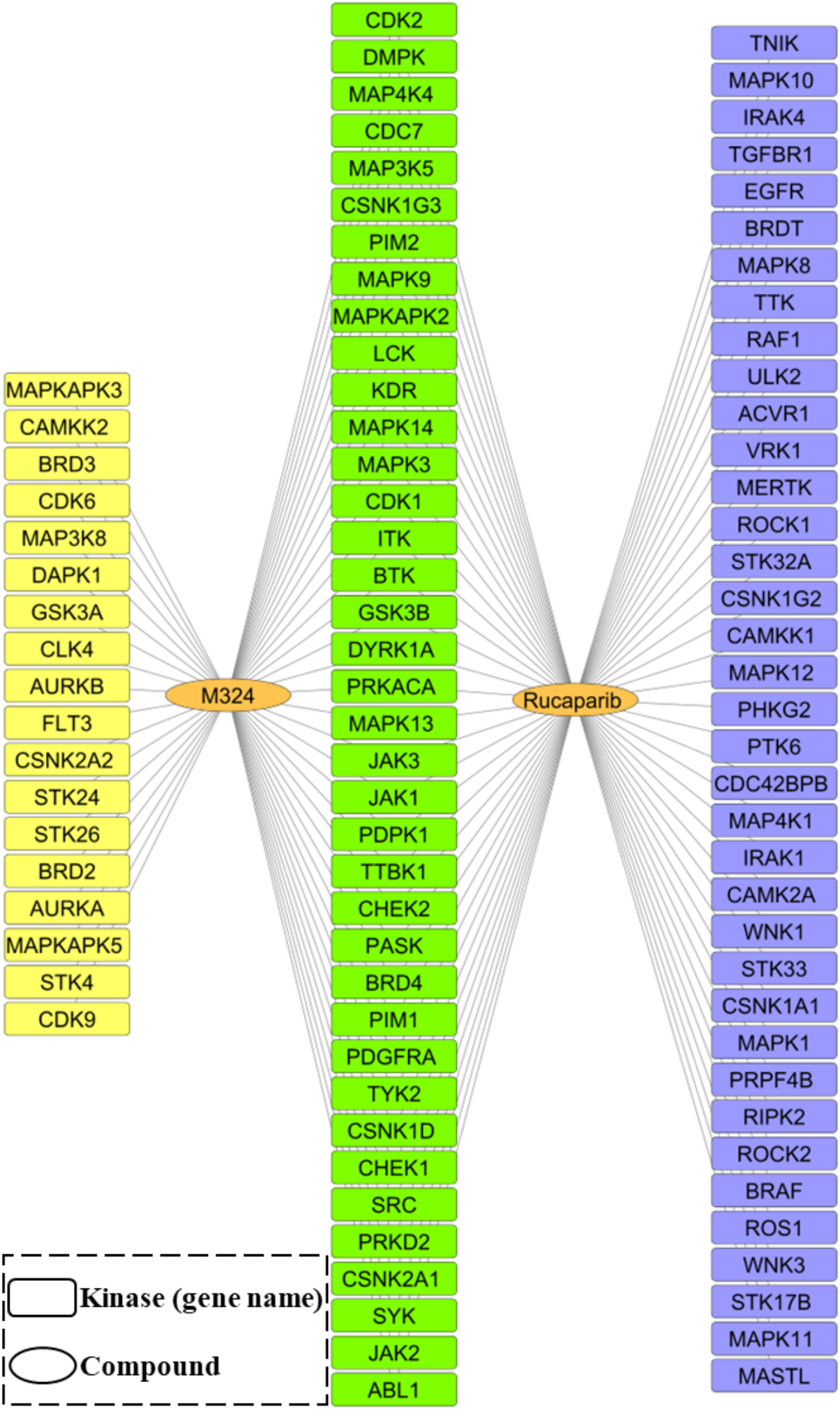
Predicted compound-kinase network. Computational predictions of the kinase profiles for M324 and rucaparib. In the network, rectangle and circular nodes represent kinases and compounds, respectively. Kinase and compound nodes are connected if the kinase is predicted as potential target for the compound through at least one of the computational methods used (see **STAR methods**). In addition, nodes coloured yellow and blue indicate that the target is predicted exclusively for M324 or for rucaparib, respectively. The kinases predicted for both compounds are highlighted in green. The gene name of each kinase target is shown inside each rectangular node.

### In vitro kinome profiling confirms differential polypharmacology between rucaparib and M324

In order to confirm the computational predictions, we decided to experimentally evaluate the kinome activity of the metabolite M324. We had previously characterized the kinome profile of rucaparib at 10 µM (Antolin et al., 2020) and identified 50 kinase off-target hits which were broadly distributed across different kinase groups, as illustrated in **Figure 4** (left). These experimental kinome results enable us to assess the performance of the computational methods: only nine out of 75 predicted kinases for rucaparib (**Table1**) were correctly predicted; all from the GalaxySagittrarius method. The other three methods (CLARITY, SEA, and PPB2) only correctly predicted PIM1 and DYRK1A, which had already been previously reported (Antolin and Mestres, 2014).

**Figure 4.**
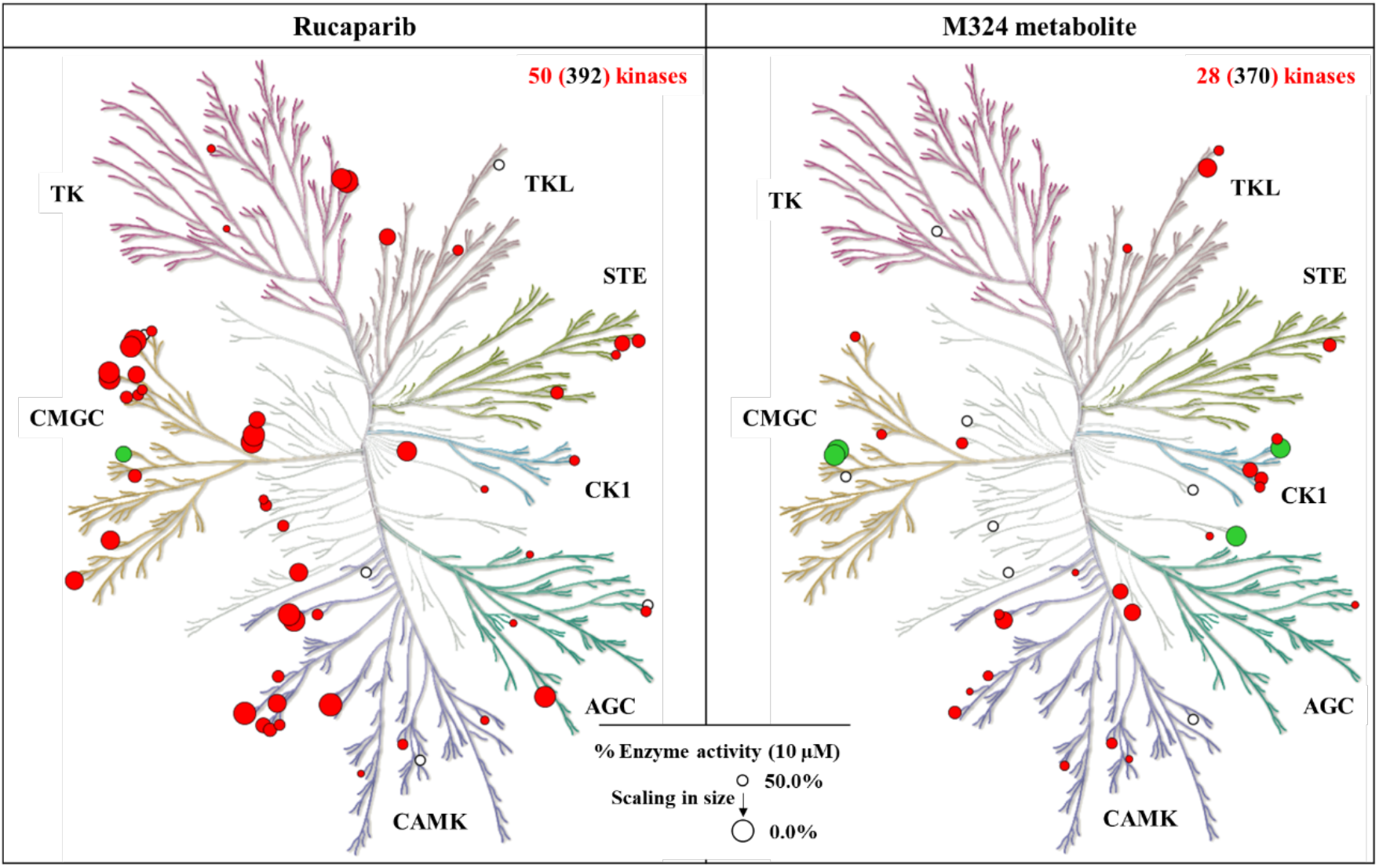
Kinome screening results for rucaparib and its major metabolite M324. Kinome trees showing the experimentally confirmed kinase hits at 10 µM concentration (red and green dots) for rucaparib (left, data derived from reference (Antolin et al., 2020)) and M324 (right, data from this work). A threshold of reducing kinase activity below 50% was applied to define a kinase hit. Kinases inhibited by more than 90% are shown in green. Circles are reversely scaled in size according to the percentage of kinase activity. White circles represent kinase hits of M324 that were not tested in rucaparib, or vice versa, due to the different assay platforms employed.

For the experimental kinome profiling of M324, we used Reaction Biology’s HotSpot platform (http://www.reactionbiology.com), that measures the inhibition of the incorporation of radiolabelled phosphate into the protein substrate to directly obtain the kinase catalytic activity (**STAR Methods**). The kinome screening results across 370 human kinases are provided in **Figure 4** (right) (and **Table S5)**. A percentage of inhibition ≥ 50% at 10 µM concentration was set as a threshold to consider potential kinase off-targets as hits. Using this cut-off, 28 kinases were identified as off-targets for M324. Although the hits were widely distributed across the kinome tree, most of the kinases were weakly inhibited (indicated by small red circles, **Figure 4**). The comparison between experimental and predicted kinase off-targets shows that nine kinases were correctly predicted for M324 (**Table 1**), six of which from GalaxySagittrarius (AURKA, PIM1, CSNK2A2, GSK3B, MAPKAPK3, and CSNK1G3). CLARITY, SEA and PPB2 correctly predicted two (AURKA and GSK3A), two (MAPKAPK3 and PASK) and three kinases (PIM1, CSNK1D, and GSK3B), respectively.

Next, we explored the overlapping kinases between rucaparib and M324. Due to the different kinase coverage between the screening platforms used, seven kinase off-targets for rucaparib were not tested for M324, and five kinases that were weakly inhibited by M324 had not been tested for rucaparib, as shown by small white circles on **Figure 4**. Excluding the assay discrepancies, eight kinases were confirmed to be inhibited by both compounds but with significant differences in the level of inhibition. For example, rucaparib showed ~ 98% inhibition against MYLK4, whereas M324 only showed ~ 70% inhibition.

Overall, the comparison between these two kinome trees (**Figure 4**) confirms the computational prediction that rucaparib and M324 display a different kinase polypharmacology profile (**Figure 3**). Specifically, rucaparib is more promiscuous than its major metabolite and strongly inhibits kinases in CMGC and CMAK groups. In contrast, M324 has a more limited number of potently inhibited kinases which are mainly distributed in the CK1 and CMGC groups.

In summary, we experimentally confirmed that rucaparib and M324 display differential polypharmacology profiles across the kinome. Due to the clinical importance of rucaparib for cancer patients, we decided to experimentally characterize these differential kinases further.

### Confirmation of submicromolar in vitro kinase activity of M324

We selected the four kinases inhibited by M324 by more than 90% (green circles on the right of **Figure 4**) – namely CSNK1E, GSK3A, GSK3B, and PLK2 – for further analysis. These four kinases belong to three different kinase families. Prior to the measurement of IC_50_ values, we tested rucaparib and M324 against these four kinases using the same catalytic inhibition assay provided by Reaction Biology (http://www.reactionbiology.com) at 1 µM concentration. As shown in **Table 2** (**Table S6**), M324 did not inhibit CSNK1E at 1 µM concentration (inactive), despite the potent inhibition detected at 10 µM (**Figure 4**; **Table S5**). M324 maintained its strong inhibition against GSK3A and PLK2 (< 50% remaining enzyme activity) and inhibited GSK3B, albeit more weakly (63.14% of remaining enzyme activity, **Table 2**). Rucaparib only strongly inhibited GSK3B, showing weak or no activity against the other three targets (**Table 2**; **Table S6**). These results further confirmed the different kinase off-target profiles between rucaparib and M324, with the latter showing much more potent inhibition against GSK3A and PLK2 than rucaparib.

**Table 2.** Confirmed kinase activity of rucaparib and M324 at 1 µM. M324 and rucaparib were screened at 1 µM against the top four kinases more strongly inhibited by M324 at 10 µM (**Figure 4** right). The tests were carried out by using a radiometric catalytic inhibition assay from Reaction Biology which directly measures the inhibition of the kinase catalytic activity. The table displays the average values of two replicate tests. The kinases more strongly inhibited that we decided to investigate further are highlighted in bold.

| Kinase group | Gene name | Enzyme activity (%) at 1μM concentration | |
| --- | --- | --- | --- |
|  |  | Rucaparib | M324 |
| CK1 | CSNK1E | 102.63 | 102.03 |
| CMGC | **GSK3A** | 74.15 | **34.35** |
| CMGC | **GSK3B** | 35.38 | **63.14** |
| Other | **PLK2** | 88.95 | **40.72** |

For completeness, we also explored the activity of the metabolite across members of the PARP enzyme family (**STAR Methods**). The metabolite shows 100% inhibition for PARP1-2 and TNKS1-2 at 10 µM concentration, and weaker inhibition for several other PARP family members, a profile that is very similar to that of rucaparib (**Table S7-8**). This similarity between both profiles across the PARP family prompted us to continue focusing on kinases to identify the main differences between the pharmacology of M324 and rucaparib.

Accordingly, we measured the *in vitro* IC_50_ values of the three kinase targets (GSK3A, GSK3B, and PLK2) that were potently inhibited by M324 or rucaparib (**Table 2**). As shown in **Figure 5a** (**Table S9**), M324 exhibits strong inhibitory activity against GSK3A (IC_50_ = 579 nM) and PLK2 (IC_50_ = 591 nM), and weakly inhibits GSK3B with single-digit micromolar activity (IC_50_ = 2.0 µM). Given the reports that M324 can permeate inside cells in animal models and reach micromolar concentrations (Murray et al., 2014), it was important to explore if these activities on isolated proteins could translate into meaningful intracellular target modulation. Therefore, we also used an *in silico* web tool (https://admetmesh.scbdd.com/) to model cell permeability in Caco-2 cells, and the results suggested that rucaparib and its metabolite have comparable permeability, with predicted values of -5.41 and -5.24 (log unit), respectively (Xiong et al., 2021). Given these results, we decided to perform follow up intracellular target engagement experiments.

**Figure 5.**
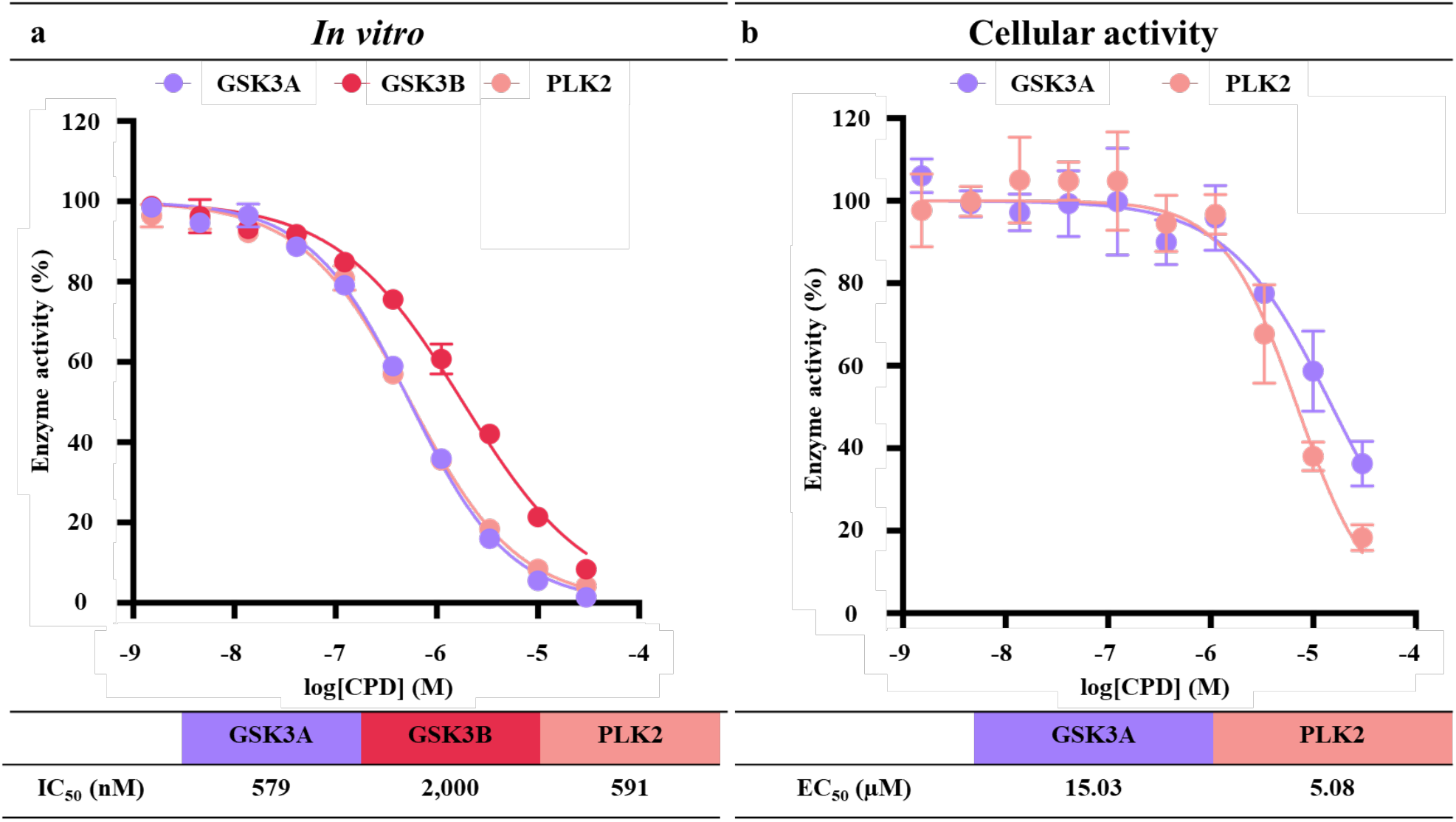
*In vitro* and intracellular kinase activity of M324. **(a)** The *in vitro* concentration-response curves and derived IC_50_ values measured in triplicate (n=3) for the three kinases that M324 more potently inhibited are depicted. **(b)** The intracellular target engagement of M324 with GSK3A (n = 2) and PLK2 (n = 4), the only two submicromolar *in vitro* kinase targets of M324, are plotted. Corresponding IC_50_ and EC_50_ values are also provided in the tables below.

### Intracellular target engagement confirms binding of M324 to PLK2 and GSK3A inside HEK293 cells

To determine the kinase activities for M324 inside cells, we used Reaction Biology’s Nano-BRET platform (http://www.reactionbiology.com) to measure intracellular target engagement for GSK3A and PLK2, the only two kinases potently inhibited by M324 with *in vitro* submicromolar activity (**Figure 5a**; **Table S9**). As shown in **Figure 5b**, the metabolite shows micromolar activity against PLK2 in HEK293 cells with EC_50_ value of 5.08 µM. In contrast, M324 shows weaker cellular activity against GSK3A (EC_50_ = 15.03 µM) (**Table S10**). Overall, these results suggest that PLK2 could be inhibited by M324 inside cells at clinically achievable micromolar concentrations.

### Rationalizing the distinct kinase inhibition profiles at the atomic level using molecular modelling

To provide a better understanding of the differential kinase off-target profiles between structurally similar rucaparib and M324, the putative binding modes between these two compounds with their kinase off-targets were modelled using the molecular docking method implemented in the MOE software (Vilar et al., 2008) (see **STAR methods** for details). Due to the absence of resolved GSK3A structures, in this study only two kinases (PLK2 and GSK3B) were analysed for rucaparib and M324.

First, we focused on PLK2, which M324 potently inhibits both *in vitro* and in cells, whereas rucaparib displays very limited kinase activity *in vitro* (around 10% inhibition at 1 µM concentration, see **Table 2** and **Table S6**). Data mining showed that only small number of PLK2 X-ray complexes were available (seven crystallographic structures are reported in the PDB (Berman et al., 2000)) and fewer than 300 IC_50_ values were deposited in the ChEMBL database ((Gaulton et al., 2017) – accessed in July 2022). Therefore, PLK2 has not been comprehensively explored in medicinal chemistry projects. Taking the resolution of the resolved PLK2 complex and the molecular size of the crystallized ligand into account, we decided to use PLK2 co-crystalized with BI 2536 (PDB ID: 4I5M) (Aubele et al., 2013) as our docking template. As shown in **Figure 6** (upper right panel), the predicted binding mode between M324 and PLK2 reveals several favourable interactions: (1) two hydrogen bonds between NHs in the M324 main scaffold and two cysteine residues (Cys67 and hinge residue Cys133) to stabilize the general binding confirmation; (2) the aromatic ring system on which fluorine is attached to form a π-π stacking interaction with residue Phe183; (3) multiple hydrogen-π interactions could be detected which were formed by residue Leu59 with the aromatic ring system of M324; (4) an electrostatic interaction. Interestingly, the negatively charged terminal carboxylic acid of M324 was favourably sandwiched between the two positively charged residues Lys57 and Arg136, as illustrated in **Figure 6** (bottom right panel).

**Figure 6.**
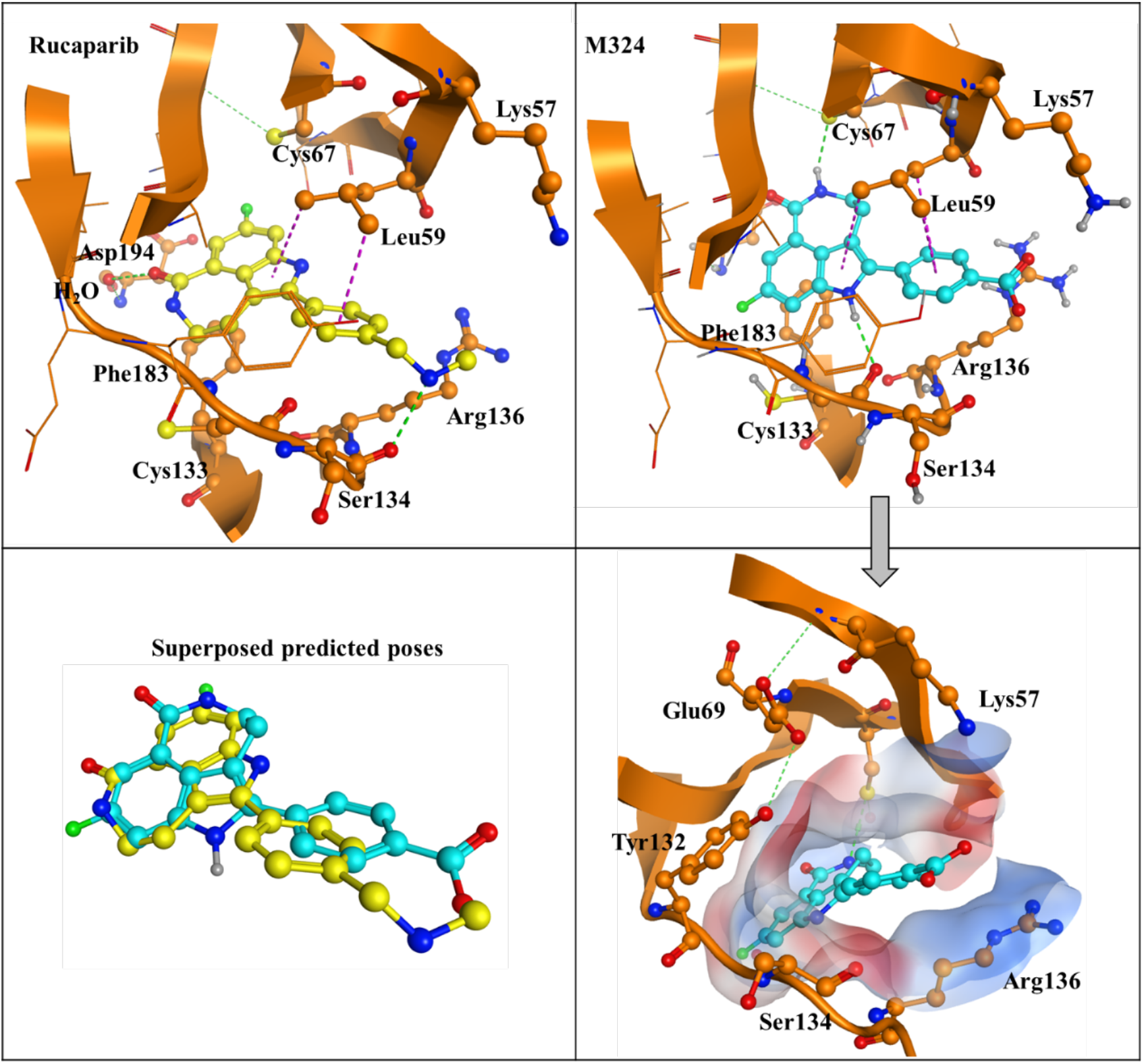
Predicted binding poses of rucaparib and M324 on PLK2 kinase. The predicted interactions of rucaparib (upper-left panel) and M324 (upper-right panel) on PLK2 kinase (PDB: 4I5M) calculated using the molecular docking environment (MOE) software are shown in the upper panel. Dashed green and magenta lines indicate hydrogen bonds and hydrogen-π interactions, respectively. The superposed predicted binding conformations of M324 and rucaparib are displayed on the bottom left panel. As it can be observed, they adopt distinct conformations, with the amine of rucaparib moving away from the positive charges of Lys57 and Arg136. The bottom right panel shows the electrostatic surface map around the binding pocket of PLK2 kinase with the binding pose of M324. The surface is colored by electrostatic potential using a continuous spectrum from blue (most positive), through white (neutral) to red (most negative). Two positively charged residues (Lys57 and Arg136) indicate the favorable electrostatic interaction with the negatively charged carboxylic acid moiety of M324.

In contrast, the analysis of interactions between rucaparib and PLK2 (upper-left corner of **Figure 6**) uncovers a different pattern. Although a hydrogen bond (with hinge residue Ser134) and two hydrogen-π interactions (with residue Leu59) could be observed, the positively charged terminal amine group in rucaparib is unfavorably positioned in the environment where two positively charged residues (Lys57 and Arg136) are located – generating a repulsive electrostatic force. Therefore, the different charges of the terminal groups of rucaparib and M324 likely explain the variations of the docking conformations between the two compounds, as shown by the superposed poses in **Figure 6** (bottom left panel). These distinct predicted interactions between rucaparib and M324 likely explain the significant difference in potency against PLK2 resulting from the structural conversion of an amine group to a carboxylic acid.

Secondly, we analyzed the putative binding modes for both compounds in GSK3B. Rucaparib exhibits superior GSK3B inhibition (~ 65 % inhibition at 1 µM concentration), nearly two-fold stronger than M324 (**Table 2**; **Table S6**), although both compounds are active. For the docking analysis, we selected the GSK3B protein structure co-crystallized with the potent ATP-competitive inhibitor alsterpaullone (PDB ID: 1Q3W) (Bertrand et al., 2003) which shares several common structural features with rucaparib. The chemical structures of alsterpaullone and rucaparib (**Figure 7**), contain a 7-membered ring with a lactone introduced in a tricyclic or tetracyclic system, which greatly contributes to their individual binding behaviors: the lactone in rucaparib interacts with two critical residues Ser904 and Gly863 in PARP1 (**Figure 1**), whereas the amide moiety in alsterpaullone forms chelated hydrogen bonds with the hinge residue Val135 in GSK3B (**Figure 7a**). However, the different position of the lactone in alsterpaullone and rucaparib implies that rucaparib might adopt a different binding confirmation to better fit into the pocket and interact with the hinge residues of GSK3B. Indeed, unlike in alsterpaullone (**Figure 7a**), in the best docking pose of rucaparib in GSK3B, its amide could not form hydrogen bonds with hinge residue Val135 (**Figure 7b)**. Instead, it showed a flipped conformation with the main scaffold containing the lactone buried into the deep ATP-binding pocket to form a hydrogen bond with residue Lys85, and the terminal moiety (the secondary amine group of rucaparib) forming two hydrogen bonds with Val135 and Pro136. The conversion of the secondary amine group to the carboxylic acid in M324 results in the loss of the hydrogen bonding network with the hinge residues, as depicted in **Figure 7b** (right). Instead, the carboxylic acid only forms one hydrogen bond with residue Tyr134. In general, the flipped lactone group could explain why both rucaparib and M324 show weaker inhibition against GSK3B in comparison with alsterpaullone. In addition, the absence of the hydrogen bond network with hinge residues observed in M324’s best predicted pose (**Figure 7b**) could further support why this metabolite exhibits lower inhibitory activity against GSK3B than its parent drug.

**Figure 7.**
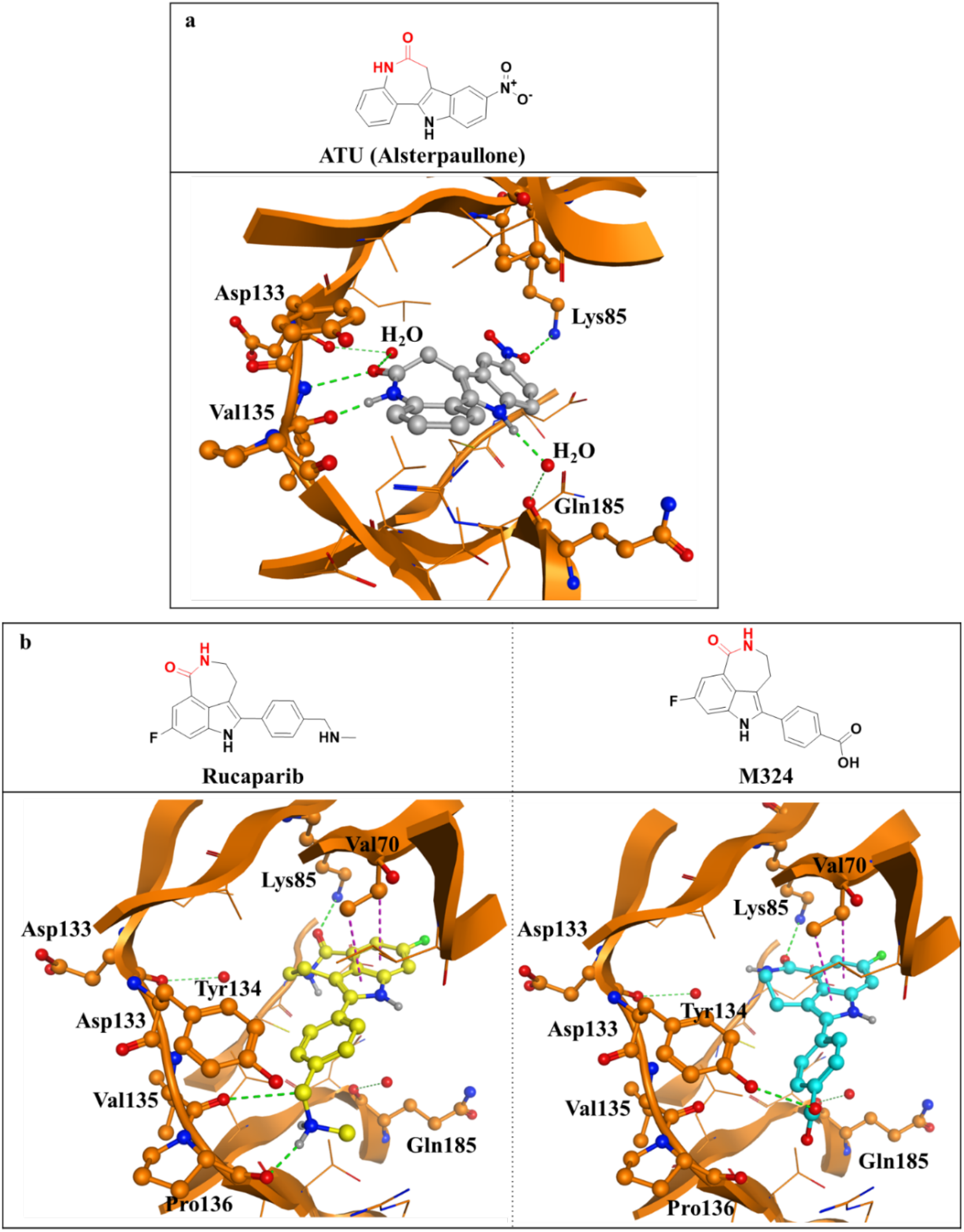
Predicted binding poses of rucaparib and M324 on GSK3B kinase. **(a)** Interactions of the ATP-competitive inhibitor alsterpaullone cocrystallized with GSK3B (PDB: 1Q3W). **(b)** Predicted binding poses and corresponding interactions of rucaparib (left) and M324 (right). As it can be observed, neither rucaparib nor M324 are able to mimic the interactions with the hinge region of the more potent GSK3B inhibitor alsterpaullone. However, rucaparib is able to form one additional hydrogen bond compared with M324, which may explain why it is more potent than its metabolite.

Taken together, for PLK2 and GSK3B, the docking analysis provides plausible explanations to rationalize the observed differential activity profiles between rucaparib and its main metabolite and help in the future design of more potent multi-target PARP-kinase inhibitors.

## Discussion

In this study, we systematically characterized the kinome profile of M324, the major metabolite of the PARP1 inhibitor rucaparib, using computational methods coupled with *in vitro* and cellular experimental validation.

The polypharmacology of drug metabolites is seldom comprehensively characterized during drug discovery and development. One of the key limitations currently hampering the study of many drug metabolites is their lack of commercial availability. Accordingly, we first synthesized the metabolite via one-step Suzuki cross-coupling reaction (**Figure 2**). The initial *in silico* prediction of kinase off-target profiles suggested that the metabolite was less promiscuous than its parent drug with 56 kinases predicted for M324 and 75 kinases for rucaparib (**Figure 3**; **Tables S1-S4**). However, both compounds had a significant number of distinct predicted kinases, suggesting that M324 could exhibit differential kinome polypharmacology compared to its parent drug (**Figure 3**). Comprehensive biochemical validation, including enzyme activity measurements at 1 and 10 µM concentration (**Figure 4**; **Table 2**; **Table S5, S6**), as well as concentration-response curves, confirmed that M324 displayed a unique kinase polypharmacology characterised by strong inhibition of GSK3A and PLK2 (IC_50_ < 600 nM) (**Figure 5a**; **Table S9**), none of which are potently inhibited by rucaparib (**Table 2**). This contrasts with the similar profile of both compounds against PARP enzymes (**Table S7, S8**). Moreover, the inhibition of these two kinases can be effectively translated into cellular activity, with M234 displaying single-digit micromolar binding to PLK2 (EC_50_ = 5.08 µM) in intracellular target engagement assays (**Figure 5b**; **Table S10**). Overall, we demonstrate for the first time that the major metabolite of the PARP1 inhibitor rucaparib can potently inhibit kinases *in vitro* and in cells. This potent inhibition of PLK2 could have implications for the clinical efficacy and safety of rucaparib and therefore warrants further investigation. Moreover, the different kinome profiles between rucaparib and its major metabolite highlights the importance of making drug metabolites more widely available and characterising them more thoroughly during drug discovery and development to comprehensively understand the therapeutic effects of drugs in the clinic and better tailor drugs to patients.

Computational methods to predict polypharmacology are in continuous development which greatly facilitates the target identification for small molecules (Cereto-Massague et al., 2015; Lavecchia and Cerchia, 2016). We have previously used computational methods relying solely on chemical similarity and reported their strengths as well as their significant limitations in comprehensively identifying polypharmacology (Antolin et al., 2020; Antolin et al., 2021a). Here, we used four different approaches (Awale and Reymond, 2019; Lounkine et al., 2012; Vidal et al., 2011; Yang et al., 2020) including GalaxySagittarius, which uses protein structure information alongside chemical information (Yang et al., 2020). GalaxySagittarius suggested dozens of kinases and showed the best recall, correctly predicting nine and six kinases for rucaparib and M324, respectively. For comparison, the other three methods provided two to three correct predictions (**Table 1**). In addition to the different algorithms employed, the prediction discrepancy observed between different computational methods could be ascribed to the varying size of the protein target database and target family coverage that they exploit for polypharmacology prediction. GalaxySagittarius uses all human proteins with crystallographic structures deposited in the PDB database with a high proportion of protein kinases (Yang et al., 2020). Therefore, we recommend the complementary use of different *in silico* approaches, particularly those using different data sources or training datasets, to provide the widest possible predictions of potential off-targets for any given compound to maximize recall.

The potent PLK2 inhibition of rucaparib’s metabolite could have clinical implications for both safety and efficacy. Unfortunately, to our knowledge, neither the free drug concentration of rucaparib or of its major metabolite have not been reported (Antolin et al., 2020). Therefore, it is difficult to draw strong conclusions but given that the reported, steady-state C_max_ concentrations of rucaparib oscillate between 2-9 µM (McCrudden et al., 2015), it is possible that M324 could significantly inhibit PLK2 in humans (EC_50_ = 5.08 µM, **Figure 5b**). Unfortunately, the physiological roles of PLK2 in health and disease have not been deeply studied (Zhang et al., 2022a). Regarding the effect of inhibiting PLK2 on human safety, a recent report suggests that the loss of function of PLK2 kinase could induce cardiac fibrosis and promote atrial fibrillation (Kunzel et al., 2021). We have recently shown that rucaparib has the highest frequency of reported cardiac adverse drug reactions, including arrhythmias (Sandhu et al., 2022) – a side effect that has not been reported with olaparib or with talazoparib. It is tempting to speculate that the unique inhibition of PLK2 by rucaparib’s metabolite could be contributing to this higher percentage of reported arrhythmias, although side effects can occur via multiple mechanisms, and it would be essential to confirm this hypothesis in clinical trials. If confirmed, these results could further support the use of alternative PARP inhibitors in patients with cardiac illnesses.

Regarding the potential effects of PLK2 inhibition in clinical efficacy, limited literature reports suggest PLK2 could be either an oncogene or a tumour suppressor depending on the context (Ou et al., 2016; Zhang et al., 2022a). Analysis of CRISPR and siRNA screens across large pan-cancer cell line panels (Dwane et al., 2021; Mitsopoulos et al., 2021; Tsherniak et al., 2017) supports this view, uncovering a few cancer cell lines that are sensitive to PLK2 knock-out, including non-small cell lung cancer and liver cancer models (**Table S11**). PLK2 inhibition could be beneficial for specific cancer subtypes and patients and detrimental for others. The potential benefits for liver cancer are particularly interesting as it has been reported that some liver cancers express cytochromes, and thus the conversion from rucaparib to M324 could occur directly inside liver cancer cells (Michael and Doherty, 2005), facilitating higher intracellular concentrations and activity of M324. Beyond oncology, PLK2 has been implicated in neurodegenerative diseases such as Parkinson’s disease that could potentially benefit from M324 (Guo et al., 2019; Weston et al., 2021). Overall, it is possible that PLK2 inhibition by M324 could have clinical implications for rucaparib’s efficacy and safety, both for cancer personalized medicine and for other indications, and this warrants further clinical investigation.

PLK2 has not been well explored in medicinal chemistry projects, as evidenced by the limited bioactivity data in ChEMBL database (version 30) (Gaulton et al., 2017). The vast majority of reported PLK2 inhibitors are derived from PLK1 projects and are the derivatives of pan-PLK inhibitor BI 2536 (Zhang et al., 2022b), largely limiting the specific study of PLK2. There is no selective, high-quality PLK2 chemical probe recommended by the Chemical Probes Portal (https://www.chemicalprobes.org/) (Antolin et al., 2022) and computational resources only identify a few selective inhibitors with modest potency (Antolin et al., 2018; Antolin et al., 2021b; Zhan et al., 2018). In this work, we show that M324 displays very high kinase selectivity for PLK2 (showing more than 80% inhibition against only seven kinases not including PLK1 and PLK3 at 10 µM concentration, **Table S5**). Moreover, we demonstrate that the metabolite inhibits PLK2 with sub-micromolar potency *in vitro* and with single-digit micromolar potency on intracellular target engagement assays (**Figure 5**). Therefore, the M324 chemotype with its tricyclic ring scaffold could be an optimal starting point to develop PLK2-selective inhibitors. In addition, our docking studies provide a plausible binding mode between M324 and PLK2 (**Figure 6**) that could guide the structure-based compound optimization towards developing much needed potent and selective PLK2 chemical probes.

Drugs often bind to more than one target (Hu and Bajorath, 2014), and there is a growing interest in rationally designing multi-target inhibitors, including the simultaneous inhibition of PARP1 and other cancer-related targets (e.g., BRD4, PI3K, EZH2, and HDACs) (Hu et al., 2022). However, the rational design of compounds active against several targets remains a non-trivial task, far from being routinely explored in drug discovery campaigns (Hu et al., 2022; Li et al., 2021). It was recently shown that there is a strong synergistic effect between PARP1 and GSK3B inhibition through drug combination studies in colorectal cancer models (Zhang et al., 2021). Our previous (Antolin et al., 2020) and current analyses suggest that rucaparib is the only approved PARP1 inhibitor exhibiting sub-micromolar inhibitory activity against GSK3B (65% of GSK3B inhibition at 1 µM concentration, **Table 2**). Therefore, rucaparib also represents a promising starting point for structural optimization to develop potent dual GSK3B-PARP inhibitors.

In conclusion, our computational and experimental analyses demonstrate that M324, the major metabolite of the PARP inhibitor rucaparib, displays a distinct kinome profile than its parent drug. This could have implications for clinical efficacy and safety and therefore warrants further clinical investigation. Therefore, these results question the preclinical study of drug action without the incorporation of abundant drug metabolites and could also open new avenues for the design of selective PLK2 inhibitors and dual PARP-kinase inhibitors. Overall, we show that drug metabolites can have different polypharmacology than their parent drugs and, therefore, they should be made more widely available and characterized more thoroughly for the benefit of personalized and precision medicine.

## METHODS

### Procedure for the preparation of metabolite M324

To a solution of 2-bromo-8-fluoro-4,5-dihydro-1H-azepino[5,4,3-cd]indol-6(3H)-one (0.16 g, 0.57 mmol) in a mixture of toluene (15 ml) and ethanol (7.5 ml), sodium carbonate (0.15 g, 1.43 mmol), 4-boronobenzoic acid (0.14 g, 0.86 mmol) and water (0.4 mL) were added sequentially. The solution was degassed with argon and Pd(Ph_3_P)_4_ (33 mg, 0.03 mmol) was added. The mixture was heated at reflux for 5 h. Then, the solution was cooled to room temperature and diluted with water (20 mL). The aqueous layer was adjusted to pH 7-8 with HCl 1N and a solid was formed. The precipitate was filtered rendering 130 mg (71%) of desired compound.

### Kinase off-target predictions using four different computational methods

In this analysis, we used four different *in silico* approaches covering various algorithms, namely: (1) the commercial target prediction software CLARITY from Chemotargets (https://www.chemotargets.com) which incorporates six independent ligand-based methods (e.g., machine learning, simplest active subgraph, etc) which were used in a consensus manner to perform the comparison of chemical structures between our compounds of interest and bioactive compounds with known binding targets (Vidal et al., 2011); (2) the public similarity ensemble approach (SEA) (https://sea.bkslab.org/), which derives macromolecule predictions based on the chemical similarity scores between the ligands (Lounkine et al., 2012); (3) the public polypharmacology browser PPB2 (https://ppb2.gdb.tools/), which features different models (e.g. fingerprint comparison, machine learning or deep learning) to predict potential targets based on bioactivity data from ChEMBL (Awale and Reymond, 2019). We used the two best performing models from PPB2: (a) nearest neighbor (NN) similarity searching with the multinomial Naive Bayes machine learning model (NB) using the ECFP4 fingerprint (**NN(ECfp4) + NB(ECfp4)**), and (b) integrating NN using shape and pharmacophore fingerprint (Xfp) with NB model of the ECFP4 fingerprint (**NN(Xfp) + NB(ECfp4)**); the predictions from both approaches were concatenated to yield the final prediction results; and (4) GalaxySagittarius (http://galaxy.seoklab.org/sagittarius) uses ligand similarity comparison to prefilter large-volume data prior to a time-consuming docking procedure (Yang et al., 2020). From the target prediction outputs of GalaxySagittarius, we selected the top 100 targets based on ranked docking scores as our results. For all four approaches, in line with our hypothesis, we only selected the targets that were human protein kinases as the final output of each method. **Table S1-S4** list the predictions obtained from these four computational methods.

### *In vitro* kinome profiling and concentration-response curves

Compounds were profiled against a kinome panel comprising 370 human kinases employing the HotSpot technology from Reaction Biology (http://www.reactionbiology.com) (Anastassiadis et al., 2011) at 10 μM concentration. Kinases more potently inhibited by M324 were further tested at 1 µM (n = 2) and/or 10-point concentration-response (n = 3). The employed *in vitro* kinase radiometric assay directly measures phosphorylation of substrate via consuming ^33^P-labelled ATP to obtain the kinase catalytic activity. Radioisotope-labelled proteins and peptides in the HotSpot assay are captured via spotting of the reaction mix on a filter membrane, whereas unreacted phosphate is washed away from the filter papers (Ma et al., 2008).

### *In vitro* PARP family profiling

All PARP family profiling assays were performed following the BPS PARP or TNKS assay kit protocols (https://bpsbioscience.com/research-areas/poly-adp-ribose-polymerase/assay-kits). The enzymatic reactions were conducted in duplicate at room temperature for 1 hour in a 96 well plate coated with histone substrate. 50 μl of reaction buffer (Tris·HCl, pH 8.0) contained NAD+, biotinylated NAD+, activated DNA, a PARP enzyme and the test compound. After enzymatic reactions, 50 μl of Streptavidin-horseradish peroxidase was added to each well and the plate was incubated at room temperature for an additional 30 min. Next, 100 μl of developer reagents were added to wells and luminescence was measured using a BioTek SynergyTM 2 microplate reader. PARP or TNKS activity assays were performed in duplicate. The luminescence data were analyzed using the computer software Graphpad Prism. In the absence of the compound, the luminescence (Lt) in each data set was defined as 100% activity. In the absence of the PARP or TNKS, the luminescence (Lb) in each data set was defined as 0% activity. The percent activity in the presence of each compound was calculated according to the following equation: % activity = [(L-Lb)/(Lt - Lb)]×100, where L= the luminescence in the presence of the compound, Lb = the luminescence in the absence of the PARP or TNKS, and Lt = the luminescence in the absence of the compound. The percent inhibition was calculated according to the following equation: % inhibition = 100 - % activity.

### Intracellular target engagement kinase assays

We used Reaction Biology’s NanoBRET platform (http://www.reactionbiology.com) which uses biophysical approaches to determine the kinase inhibitor occupancy of a ligand in intact living cells using BRET and an optimized cell-permeable kinase tracer. The specificity of the BRET signal is dictated by the placement of NanoLuc on the specific kinase and transfected into HEK293 cells, which were established from primary embryonic human kidney.

Human embryonic kidney (HEK293) cells were purchased from ATCC. FuGENER HD Transfection Reagent, KinaseNanoLuc® fusion plasmids, Transfection Carrier DNA, NanoBRET™ Tracer and dilution buffer, NanoBRET™ Nano-Glo® Substrate, Extracellular NanoLuc® Inhibitor were from Promega. AT7519 and BI-2536 were used as positive controls for the determination cellular activities of GSK3A and PLK2, respectively.

HEK293 cells were transiently transfected with the KinaseNanoLuc® Fusion Vector DNA using FuGENER HD Transfection Reagent. Test compounds were dispensed into 384 well assay plate using an Echo 550 acoustic dispenser (Labcyte Inc, Sunnyvale, CA). Transfected cells were harvested and mixed with NanoBRET™ Tracer Reagent and dispensed into 384 well plates and incubated at 37C in 5% CO_2_ cell culture incubator for 1 hour. The NanoBRET™ Nano-Glo® Substrate plus Extracellular NanoLuc® Inhibitor Solution were added into the wells of the assay plate and incubated for 2–3 minutes at room temperature. The donor emission wavelength (460 nm) and acceptor emission wavelength (600 nm) were measured in an EnVision plate reader. The BRET Ratio was calculated using the equation: BRET Ratio = [(Acceptor sample ÷ Donor sample) – (Acceptor no-tracer control ÷ Donor no-tracer control)].

### Docking studies

To explore the molecular interactions of M324 and rucaparib with their stronger kinase hits we used the docking method available in MOE 2020.09 (https://www.chemcomp.com/Products.htm) (Vilar et al., 2008). From the three kinases more potently inhibited by M324 *in vitro*, only two – GSK3B and PLK2 – had a crystal structure deposited in the PDB and, therefore, they are the focus of this study. We choose the X-ray complex with the closest molecular size or most similar scaffold to rucaparib to increase the accuracy of the modelling: (1) the crystallographic structure of PLK2 in complex with BI 2536 (PDB ID: 4I5M), an ATP-competitive inhibitor, was used to predict the interactions of rucaparib and M324 on PLK2 kinase; (2) the X-ray complex co-crystallized with alsterpaullone (PDB ID: 1Q3W), which contains an amide-bearing tetracyclic scaffold, was selected for GSK3B. Prior to the docking studies, crystal structures were corrected for missing atoms and bonds, and protonated under physiological condition using standard methods available in MOE. Water molecules were only removed provided the absence of water-mediated interactions in original structures. The co-crystalized ligands were exploited to define the active site and the general default parameters for docking were applied.

## Acknowledgements

A.A.A. was primarily supported by an ICR Fellowship and was formerly supported by a Wellcome Trust Sir Henry Wellcome Postdoctoral Fellowship (204735/Z/16/Z). C.S and A.L thank CSIC (201980E011) and Catalan Government (2017SGR 1604) for their support. The authors thank many colleagues and collaborators for helpful discussions and valuable input into the preparation of this manuscript.

## Author contributions

A.A.A., H.H., C.S., and A.L. designed the research. H.H. and A.A.A. performed the computational research and managed the experimental biology work. C.S and A.L. designed and performed the chemical synthesis. A.A.A., H.H., C.S. and A.L. conducted data analysis and interpretation. A.A.A. and H.H. wrote the manuscript with contributions from C.S. and A.L.

## References

Anastassiadis, T., Deacon, S.W., Devarajan, K., Ma, H., and Peterson, J.R. (2011). Comprehensive assay of kinase catalytic activity reveals features of kinase inhibitor selectivity. Nat. Biotechnol. 29, 1039–1045. https://doi.org/10.1038/nbt.2017.

Anighoro, A., Bajorath, J., and Rastelli, G. (2014). Polypharmacology: challenges and opportunities in drug discovery. J. Med. Chem. 57, 7874–7887. https://doi.org/10.1021/jm5006463.

Antolin, A.A., Ameratunga, M., Banerji, U., Clarke, P.A., Workman, P., and Al-Lazikani, B. (2020). The kinase polypharmacology landscape of clinical PARP inhibitors. Sci. Rep. 10, 2585. https://doi.org/10.1038/s41598-020-59074-4.

Antolin, A.A., Clarke, P.A., Collins, I., Workman, P., and Al-Lazikani, B. (2021a). Evolution of kinase polypharmacology across HSP90 drug discovery. Cell Chem. Biol. 28, 1433–1445 e1433. https://doi.org/10.1016/j.chembiol.2021.05.004.

Antolin, A.A., and Mestres, J. (2014). Linking off-target kinase pharmacology to the differential cellular effects observed among PARP inhibitors. Oncotarget 5, 3023–3028. https://doi.org/10.18632/oncotarget.1814.

Antolin, A.A., Tym, J.E., Komianou, A., Collins, I., Workman, P., and Al-Lazikani, B. (2018). Objective, quantitative, data-driven assessment of chemical probes. Cell Chem. Biol. 25, 194–205 e195. https://doi.org/10.1016/j.chembiol.2017.11.004.

Antolin, A.A., Workman, P., and Al-Lazikani, B. (2021b). Public resources for chemical probes: the journey so far and the road ahead. Future Med. Chem. 13, 731–747. https://doi.org/10.4155/fmc-2019-0231.

Antolin, A.A., Sanfelice, D., Crisp, A., Villasclaras Fernandez, E., Mica, I.L., Chen, Y., Collins, I., Edwards, A., Müller, S., Al-Lazikani, B., Workman, P. (2022) The Chemical Probes Portal: an expert review-based public resource to empower chemical probe assessment, selection and use. Nucleic Acids Res. In press. https://doi.org/10.1093/nar/gkac909.

Aubele, D.L., Hom, R.K., Adler, M., Galemmo, R.A., Jr., Bowers, S., Truong, A.P., Pan, H., Beroza, P., Neitz, R.J., Yao, N., et al. (2013). Selective and brain-permeable polo-like kinase-2 (Plk-2) inhibitors that reduce alpha-synuclein phosphorylation in rat brain. ChemMedChem 8, 1295–1313. https://doi.org/10.1002/cmdc.201300166.

Awale, M., and Reymond, J.L. (2019). Polypharmacology Browser PPB2: target prediction combining nearest neighbors with machine learning. J. Chem. Inf. Model. 59, 10–17. https://doi.org/10.1021/acs.jcim.8b00524.

Baillie, T.A., and Rettie, A.E. (2011). Role of biotransformation in drug-induced toxicity: influence of intra- and inter-species differences in drug metabolism. Drug Metab. Pharmacokinet. 26, 15–29. https://doi.org/10.2133/dmpk.dmpk-10-rv-089.

Bender, R.P., Lindsey, R.H., Jr., Burden, D.A., and Osheroff, N. (2004). N-acetyl-p-benzoquinone imine, the toxic metabolite of acetaminophen, is a topoisomerase II poison. Biochemistry 43, 3731–3739. https://doi.org/10.1021/bi036107r.

Berman, H.M., Westbrook, J., Feng, Z., Gilliland, G., Bhat, T.N., Weissig, H., Shindyalov, I.N., and Bourne, P.E. (2000). The protein data bank. Nucleic Acids Res. 28, 235–242. https://doi.org/10.1093/nar/28.1.235.

Bertrand, J.A., Thieffine, S., Vulpetti, A., Cristiani, C., Valsasina, B., Knapp, S., Kalisz, H.M., and Flocco, M. (2003). Structural characterization of the GSK-3beta active site using selective and non-selective ATP-mimetic inhibitors. J. Mol. Biol. 333, 393–407. https://doi.org/10.1016/j.jmb.2003.08.031.

Bolognesi, M.L. (2019). Harnessing polypharmacology with medicinal chemistry. ACS Med. Chem. Lett. 10, 273–275. https://doi.org/10.1021/acsmedchemlett.9b00039.

Cereto-Massague, A., Ojeda, M.J., Valls, C., Mulero, M., Pujadas, G., and Garcia-Vallve, S. (2015). Tools for in silico target fishing. Methods 71, 98–103. https://doi.org/10.1016/j.ymeth.2014.09.006.

Dwane, L., Behan, F.M., Goncalves, E., Lightfoot, H., Yang, W., van der Meer, D., Shepherd, R., Pignatelli, M., Iorio, F., and Garnett, M.J. (2021). Project score database: a resource for investigating cancer cell dependencies and prioritizing therapeutic targets. Nucleic Acids Res. 49, D1365–D1372. https://doi.org/10.1093/nar/gkaa882.

Eid, S., Turk, S., Volkamer, A., Rippmann, F., and Fulle, S. (2017). KinMap: a web-based tool for interactive navigation through human kinome data. BMC Bioinf. 18, 16. https://doi.org/10.1186/s12859-016-1433-7.

Ferraris, D.V. (2010). Evolution of poly(ADP-ribose) polymerase-1 (PARP-1) inhibitors. From concept to clinic. J. Med. Chem. 53, 4561–4584. https://doi.org/10.1021/jm100012m.

Fura, A. (2006). Role of pharmacologically active metabolites in drug discovery and development. Drug Discov. Today 11, 133–142. https://doi.org/10.1016/S1359-6446(05)03681-0.

Gaulton, A., Hersey, A., Nowotka, M., Bento, A.P., Chambers, J., Mendez, D., Mutowo, P., Atkinson, F., Bellis, L.J., Cibrian-Uhalte, E., et al. (2017). The ChEMBL database in 2017. Nucleic Acids Res. 45, D945–D954. https://doi.org/10.1093/nar/gkw1074.

Guengerich, F.P. (2001). Common and uncommon cytochrome P450 reactions related to metabolism and chemical toxicity. Chem. Res. Toxicol. 14, 611–650. https://doi.org/10.1021/tx0002583.

Guo, C., Zhu, J., Wang, J., Duan, J., Ma, S., Yin, Y., Quan, W., Zhang, W., Guan, Y., Ding, Y., et al. (2019). Neuroprotective effects of protocatechuic aldehyde through PLK2/p-GSK3beta/Nrf2 signaling pathway in both in vivo and in vitro models of Parkinson’s disease. Aging (Albany NY) 11, 9424–9441. https://doi.org/10.18632/aging.102394.

Hopkins, A.L. (2008). Network pharmacology: the next paradigm in drug discovery. Nat. Chem. Biol. 4, 682–690. https://doi.org/10.1038/nchembio.118.

Hu, X., Zhang, J., Zhang, Y., Jiao, F., Wang, J., Chen, H., Ouyang, L., and Wang, Y. (2022). Dual-target inhibitors of poly (ADP-ribose) polymerase-1 for cancer therapy: Advances, challenges, and opportunities. Eur J Med Chem 230, 114094. https://doi.org/10.1016/j.ejmech.2021.114094.

Hu, Y., and Bajorath, J. (2014). Monitoring drug promiscuity over time. F1000Res. 3, 218. https://doi.org/10.12688/f1000research.5250.2.

Kabir, A., and Muth, A. (2022). Polypharmacology: The science of multi-targeting molecules. Pharmacol. Res. 176, 106055. https://doi.org/10.1016/j.phrs.2021.106055.

Knezevic, C.E., Wright, G., Rix, L.L.R., Kim, W., Kuenzi, B.M., Luo, Y., Watters, J.M., Koomen, J.M., Haura, E.B., Monteiro, A.N., et al. (2016). Proteome-wide profiling of clinical PARP inhibitors reveals compound-specific secondary targets. Cell Chem. Biol. 23, 1490–1503. https://doi.org/10.1016/j.chembiol.2016.10.011.

Kunzel, S.R., Hoffmann, M., Weber, S., Kunzel, K., Kammerer, S., Gunscht, M., Klapproth, E., Rausch, J.S.E., Sadek, M.S., Kolanowski, T., et al. (2021). Diminished PLK2 induces cardiac fibrosis and promotes atrial fibrillation. Circ. Res. 129, 804–820. https://doi.org/10.1161/CIRCRESAHA.121.319425.

LaFargue, C.J., Dal Molin, G.Z., Sood, A.K., and Coleman, R.L. (2019). Exploring and comparing adverse events between PARP inhibitors. Lancet Oncol. 20, e15–e28. https://doi.org/10.1016/S1470-2045(18)30786-1.

Lavecchia, A., and Cerchia, C. (2016). In silico methods to address polypharmacology: current status, applications and future perspectives. Drug Discov. Today 21, 288–298. https://doi.org/10.1016/j.drudis.2015.12.007.

Li, X., Li, X., Liu, F., Li, S., and Shi, D. (2021). Rational multitargeted drug design strategy from the perspective of a medicinal chemist. J. Med. Chem. 64, 10581–10605. https://doi.org/10.1021/acs.jmedchem.1c00683.

Liao, M., Watkins, S., Nash, E., Isaacson, J., Etter, J., Beltman, J., Fan, R., Shen, L., Mutlib, A., Kemeny, V., et al. (2020). Evaluation of absorption, distribution, metabolism, and excretion of [(14)C]-rucaparib, a poly(ADP-ribose) polymerase inhibitor, in patients with advanced solid tumors. Invest. New Drugs 38, 765–775. https://doi.org/10.1007/s10637-019-00815-2.

Lounkine, E., Keiser, M.J., Whitebread, S., Mikhailov, D., Hamon, J., Jenkins, J.L., Lavan, P., Weber, E., Doak, A.K., Cote, S., et al. (2012). Large-scale prediction and testing of drug activity on side-effect targets. Nature 486, 361–367. https://doi.org/10.1038/nature11159.

Ma, H., Deacon, S., and Horiuchi, K. (2008). The challenge of selecting protein kinase assays for lead discovery optimization. Expert Opin. Drug Discov. 3, 607–621. https://doi.org/10.1517/17460441.3.6.607.

Mateo, J., Lord, C.J., Serra, V., Tutt, A., Balmana, J., Castroviejo-Bermejo, M., Cruz, C., Oaknin, A., Kaye, S.B., and de Bono, J.S. (2019). A decade of clinical development of PARP inhibitors in perspective. Ann. Oncol. 30, 1437–1447. https://doi.org/10.1093/annonc/mdz192.

McCrudden, C.M., O’Rourke, M.G., Cherry, K.E., Yuen, H.F., O’Rourke, D., Babur, M., Telfer, B.A., Thomas, H.D., Keane, P., Nambirajan, T., et al. (2015). Vasoactivity of rucaparib, a PARP-1 inhibitor, is a complex process that involves myosin light chain kinase, P2 receptors, and PARP itself. PLoS One 10, e0118187. https://doi.org/10.1371/journal.pone.0118187.

Michael, M., and Doherty, M.M. (2005). Tumoral drug metabolism: overview and its implications for cancer therapy. J. Clin. Oncol. 23, 205–229. https://doi.org/10.1200/JCO.2005.02.120.

Mitsopoulos, C., Di Micco, P., Fernandez, E.V., Dolciami, D., Holt, E., Mica, I.L., Coker, E.A., Tym, J.E., Campbell, J., Che, K.H., et al. (2021). canSAR: update to the cancer translational research and drug discovery knowledgebase. Nucleic Acids Res. 49, D1074–D1082. https://doi.org/10.1093/nar/gkaa1059.

Murray, J., Thomas, H., Berry, P., Kyle, S., Patterson, M., Jones, C., Los, G., Hostomsky, Z., Plummer, E.R., Boddy, A.V., and Curtin, N.J. (2014). Tumour cell retention of rucaparib, sustained PARP inhibition and efficacy of weekly as well as daily schedules. Br. J. Cancer 110, 1977–1984. https://doi.org/10.1038/bjc.2014.91.

Ou, B., Zhao, J., Guan, S., Wangpu, X., Zhu, C., Zong, Y., Ma, J., Sun, J., Zheng, M., Feng, H., and Lu, A. (2016). PLK2 promotes tumor growth and inhibits apoptosis by targeting Fbxw7/Cyclin E in colorectal cancer. Cancer Lett. 380, 457–466. https://doi.org/10.1016/j.canlet.2016.07.004.

Rose, M., Burgess, J.T., O’Byrne, K., Richard, D.J., and Bolderson, E. (2020). PARP Inhibitors: clinical relevance, mechanisms of action and tumor resistance. Front Cell Dev. Biol. 8, 564601. https://doi.org/10.3389/fcell.2020.564601.

Sandhu, D., Antolin, A.A., Cox, A.R., and Jones, A.M. (2022). Identification of different side effects between PARP inhibitors and their polypharmacological multi-target rationale. Br. J. Clin. Pharmacol. 88, 742–752. https://doi.org/10.1111/bcp.15015.

Sinha, S., Molla, S., and Kundu, C.N. (2021). PARP1-modulated chromatin remodeling is a new target for cancer treatment. Med. Oncol. 38, 118. https://doi.org/10.1007/s12032-021-01570-2.

Thompson, R.A., Isin, E.M., Ogese, M.O., Mettetal, J.T., and Williams, D.P. (2016). Reactive metabolites: current and emerging risk and hazard assessments. Chem. Res. Toxicol. 29, 505–533. https://doi.org/10.1021/acs.chemrestox.5b00410.

Thorsell, A.G., Ekblad, T., Karlberg, T., Low, M., Pinto, A.F., Tresaugues, L., Moche, M., Cohen, M.S., and Schuler, H. (2017). Structural basis for potency and promiscuity in Poly(ADP-ribose) polymerase (PARP) and tankyrase inhibitors. J. Med. Chem. 60, 1262–1271. https://doi.org/10.1021/acs.jmedchem.6b00990.

Tsherniak, A., Vazquez, F., Montgomery, P.G., Weir, B.A., Kryukov, G., Cowley, G.S., Gill, S., Harrington, W.F., Pantel, S., Krill-Burger, J.M., et al. (2017). Defining a cancer dependency map. Cell 170, 564–576 e516. https://doi.org/10.1016/j.cell.2017.06.010.

Vidal, D., Garcia-Serna, R., and Mestres, J. (2011). Ligand-based approaches to in silico pharmacology. Methods Mol. Biol. 672, 489–502. https://doi.org/10.1007/978-1-60761-839-3_19.

Vilar, S., Cozza, G., and Moro, S. (2008). Medicinal chemistry and the molecular operating environment (MOE): application of QSAR and molecular docking to drug discovery. Curr. Top. Med. Chem. 8, 1555–1572. https://doi.org/10.2174/156802608786786624.

Walther, R., Rautio, J., and Zelikin, A.N. (2017). Prodrugs in medicinal chemistry and enzyme prodrug therapies. Adv. Drug Deliv. Rev. 118, 65–77. https://doi.org/10.1016/j.addr.2017.06.013.

Weston, L.J., Stackhouse, T.L., Spinelli, K.J., Boutros, S.W., Rose, E.P., Osterberg, V.R., Luk, K.C., Raber, J., Weissman, T.A., and Unni, V.K. (2021). Genetic deletion of Polo-like kinase 2 reduces alpha-synuclein serine-129 phosphorylation in presynaptic terminals but not Lewy bodies. J. Biol. Chem. 296, 100273. https://doi.org/10.1016/j.jbc.2021.100273.

Xiong, G., Wu, Z., Yi, J., Fu, L., Yang, Z., Hsieh, C., Yin, M., Zeng, X., Wu, C., Lu, A., et al. (2021). ADMETlab 2.0: an integrated online platform for accurate and comprehensive predictions of ADMET properties. Nucleic Acids Res. 49, W5–W14. https://doi.org/10.1093/nar/gkab255.

Yang, J., Kwon, S., Bae, S.H., Park, K.M., Yoon, C., Lee, J.H., and Seok, C. (2020). GalaxySagittarius: structure- and similarity-based prediction of protein targets for druglike compounds. J. Chem. Inf. Model. 60, 3246–3254. https://doi.org/10.1021/acs.jcim.0c00104.

Zhan, M.M., Yang, Y., Luo, J., Zhang, X.X., Xiao, X., Li, S., Cheng, K., Xie, Z., Tu, Z., and Liao, C. (2018). Design, synthesis, and biological evaluation of novel highly selective polo-like kinase 2 inhibitors based on the tetrahydropteridin chemical scaffold. Eur J Med Chem 143, 724–731. https://doi.org/10.1016/j.ejmech.2017.11.058.

Zhang, C., Ni, C., and Lu, H. (2022a). Polo-like kinase 2: from principle to practice. Front. Oncol. 12, 956225. https://doi.org/10.3389/fonc.2022.956225.

Zhang, J., Zhang, L., Wang, J., Ouyang, L., and Wang, Y. (2022b). Polo-like kinase 1 inhibitors in human cancer therapy: development and therapeutic potential. J. Med. Chem. 65, 10133–10160. https://doi.org/10.1021/acs.jmedchem.2c00614.

Zhang, N., Tian, Y.N., Zhou, L.N., Li, M.Z., Chen, H.D., Song, S.S., Huan, X.J., Bao, X.B., Zhang, A., Miao, Z.H., and He, J.X. (2021). Glycogen synthase kinase 3beta inhibition synergizes with PARP inhibitors through the induction of homologous recombination deficiency in colorectal cancer. Cell Death Dis. 12, 183. https://doi.org/10.1038/s41419-021-03475-4.

Zhang, Z., and Tang, W. (2018). Drug metabolism in drug discovery and development. Acta Pharm. Sin. B 8, 721–732. https://doi.org/10.1016/j.apsb.2018.04.003.

