## Supplementary Figures 1-2 for "Identification of differential polypharmacology between the PARP inhibitor rucaparib and its major metabolite"

**Figure S1. HPLC/MS Analysis details**. Column: ZORBAX Extend-C18 3.5 µm 2.1 x 50 mm. Temperature: 30ºC. A: H2O + 0.1% DEA + 40mM AcONH4 (pH 9.4) B: can. Inyection volume: 5µl. Conditions of the MS detector LTQ XL ESI ‐ ion trap (Thermo Scientific): Positive and negative mode; m/z 50 ‐ 2000; Capillary temperature: 300ºC; Sheath gas flow (N2): 60; Auxiliary gas flow (N2): 10; Sweep gas flow (N2): 10; Damping gas: He; Capillary voltage: 32V positive mode/ ‐ 24V negative mode; Source voltage: 4.5kV; Source current: 100 uA; Tube Lens: 70V positive mode/ ‐ 103.20V negative mode; Full micro scans: 1; Full Max Ion time: 10ms.

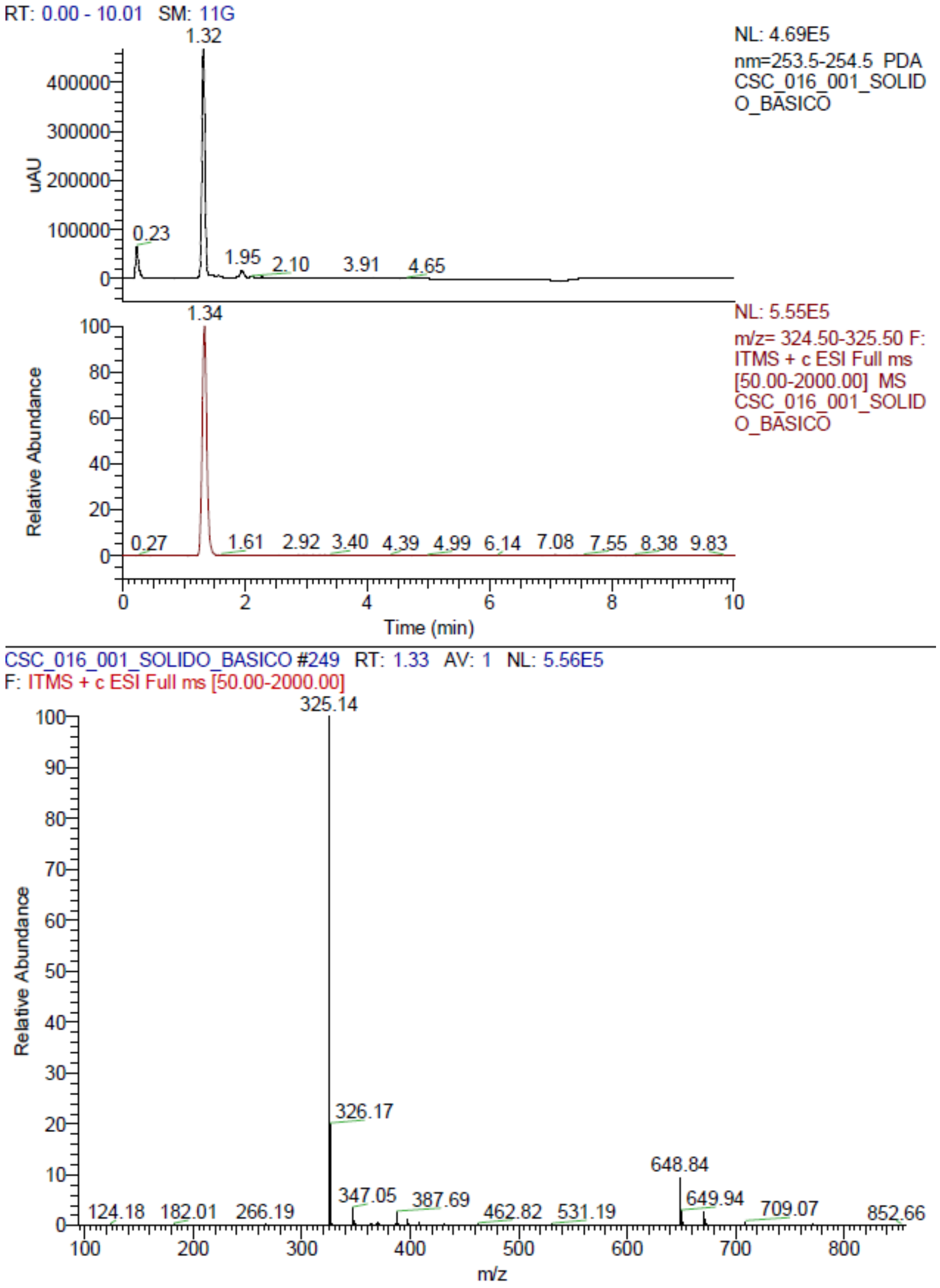

**Figure S2. NMR Analysis details**. a) Proton-NMR (including enlarged subsection in the center); b) Carbon-NMR; c) Carbon-NMR (enlarged subsection); d) gDQCOSY; e) HSQCAD; f) gHMBCAD.

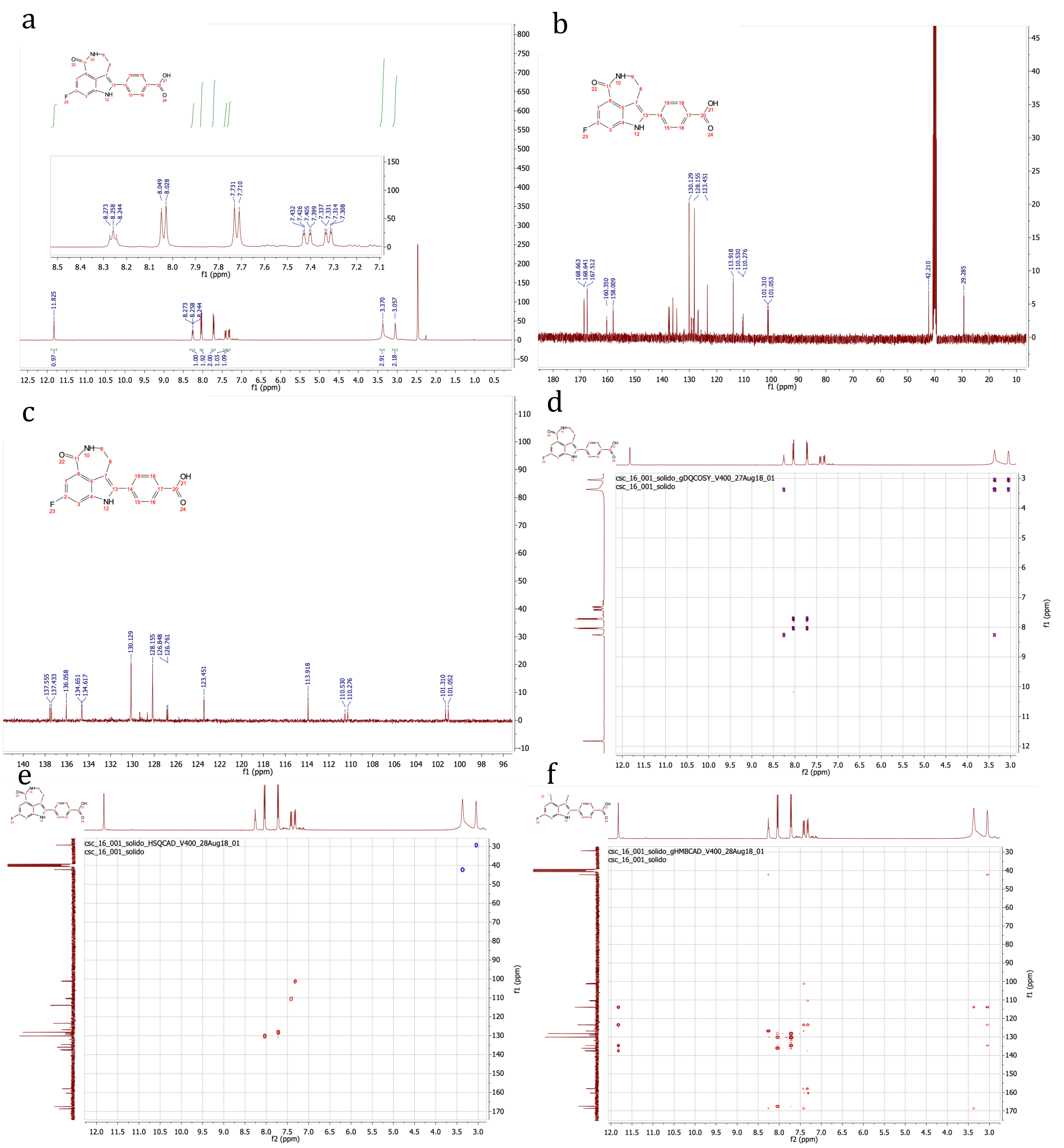
